# Automatic discrimination of species within the *Enterobacter cloacae* complex using MALDI-TOF Mass Spectrometry and supervised algorithms

**DOI:** 10.1101/2021.11.02.467040

**Authors:** Ana Candela, Alejandro Guerrero-López, Miriam Mateos, Alicia Gómez-Asenjo, Manuel J. Arroyo, Marta Hernandez-García, Rosa del Campo, Emilia Cercenado, Aline Cuénod, Gema Méndez, Luis Mancera, Juan de Dios Caballero, Laura Martínez-García, Desirée Gijón, María Isabel Morosini, Patricia Ruiz-Garbajosa, Adrian Egli, Rafael Cantón, Patricia Muñoz, David Rodríguez-Temporal, Belén Rodríguez-Sánchez

## Abstract

The *Enterobacter cloacae* complex (ECC) encompasses heterogeneous clusters of species that have been associated with nosocomial outbreaks. These species may host different acquired antimicrobial resistance and virulence mechanisms and their identification are challenging. This study aims to develop predictive models based on MALDI-TOF MS spectral profiles and machine learning for species-level identification.

A total of 198 ECC and 116 *K. aerogenes* clinical isolates from the University Hospital Ramón y Cajal (Spain) and the University Hospital Basel (Switzerland) were included. The capability of the proposed method to differentiate the most common ECC species (*E. asburiae, E. kobei, E. hormaechei, E. roggenkampii, E. ludwigii, E. bugandensis*) and *K. aerogenes* was demonstrated by applying unsupervised hierarchical clustering with PCA pre-processing. We observed a distinctive clustering of *E. hormaechei* and *K. aerogenes* and a clear trend for the rest of the ECC species to be differentiated over the development dataset. Thus, we developed supervised, non-linear predictive models (Support Vector Machine with Radial Basis Function and Random Forest). The external validation of these models with protein spectra from the two participating hospitals yielded 100% correct species-level assignment for *E. asburiae, E. kobei*, and *E. roggenkampii* and between 91.2% and 98.0% for the remaining ECC species. Similar results were obtained with the MSI database developed recently (https://msi.happy-dev.fr/) except in the case of *E. hormaechei*, which was more accurately identified by Random Forest.

In short, MALDI-TOF MS combined with machine learning demonstrated to be a rapid and accurate method for the differentiation of ECC species.

## INTRODUCTION

*Enterobacter* is a facultative anaerobic Gram-negative genus that can be found as a natural commensal in the gut microbiome of mammals (1). Several species have been associated with nosocomial outbreaks causing urinary tract infection, skin and soft tissue infection, pneumonia, and bacteremia (2, 3). *Enterobacter cloacae* complex (ECC) is of particular clinical interest. This group is composed of 13 heterogenic genetic clusters according to *hsp60* gene sequencing: *E. asburiae* (cluster I), *E. kobei* (cluster II), *E. hormaechei* subsp. *hoffmannii* (cluster III), *E. roggenkampii* (cluster IV), *E. ludwigii* (cluster V), *E. hormaechei* subsp. *oharae*, and subsp. *xiangfangensis* (cluster VI), *E. hormaechei* subsp. *hormaechei* (cluster VII), *E. hormaechei* subsp. *steigerwaltii* (cluster VIII), *E. bugandensis* (cluster IX), *E. nimipressuralis* (cluster X), *E. cloacae* subsp. *cloacae* (cluster XI), *E. cloacae* subsp. *dissolvens* (cluster XII), and a heterogeneous group of *E. cloacae* sequences are considered as cluster XIII. However, the taxonomy of this genus is still under debate (4, 5). In fact, *Enterobacter aerogenes* has been recently reclassified into the *Klebsiella* genus as *K. aerogenes* (6). A more comprehensive study based on whole-genome sequencing (WGS) data from ECC isolates yielded a redistribution of the species defined by *hsp60* sequencing (5) into 22 clades (7) and allowed the characterization of new ECC species (8).

Discrimination of the ECC at the species level is usually performed by sequenc-based methods. The most commonly targeted gene is *hsp60*, although multi-locus sequence typing (MLST) and WGS have also been applied (5, 9, 10). Sequence-based diagnostic methods techniques are laborious and require specific equipment. Therefore, new emerging techniques such as Matrix Assisted Laser Desorption/Ionization Time-of-flight Mass Spectrometry (MALDI-TOF MS) have been proposed as an alternative to sequence-based methods. MALDI-TOF MS has shown to be an excellent methodology for bacterial identification. It can easily identify *E. cloacae* complex isolates but it showed low discrimination power for the species in this group when using standard analysis and commercial databases with low resolution (11, 12).

This study aimed to develop and validate prediction models for the automatic species differentiation within the ECC using MALDI-TOF MS and supervised learning algorithms. This task is important because of the diverse implications of ECC species in human pathologies and their involvement in nosocomial outbreaks (4). Besides, *E. hormaechei*, the most encountered ECC species in the clinical settings, has been correlated with the enhanced acquisition of antimicrobial resistance mechanisms and the expression of virulence factors (13, 14). To achieve this goal, two steps were conducted in this study. First, we performed an unsupervised clustering to determine the feasibility of MALDI-TOF MS data for the ECC species identification. Second, we applied a supervised machine learning algorithm with isolates from University Hospital Ramón y Cajal, Madrid -Madrid, Spain - and validated our findings with different ECC isolates from the same hospital and from the University Hospital of Basel, Switzerland.

## MATERIALS AND METHODS

### Bacterial isolates

Overall, we analyzed 198 clinical isolates belonging to the ECC and nine 116 *K. aerogenes* (formerly *E. aerogenes*). Among them, 164 ECC and 9 *K. aerogenes* were collected in a surveillance study of antimicrobial resistance in the Hospital Universitario Ramón y Cajal -UHRC- (Madrid, Spain) between 2005 and 2018 and identified by partial sequencing of the *hsp60* gene (15). We collected the remaining isolates (34 ECC and 107 *K. aerogenes*) at the University Hospital Basel (UHB; Basel, Switzerland) between 2016 and 2021 and identified the isolates by whole genome sequencing (WGS) using KmerFinder 3.2 (16–18). MALDI-TOF MS spectral profiles of these isolates were obtained in Basel and submitted to the Hospital General Universitario Gregorio Marañón (Madrid, Spain) for further analysis.

All isolates from UHRC were incubated overnight at 37°C and metabolically activated after three subcultures on Columbia Blood Agar (bioMérieux, Marcy l’Etoile, France) before their analysis with MALDI-TOF MS at the Hospital General Universitario Gregorio Marañón.

### Spectra acquisition using MALDI-TOF MS

We identified the isolates using the MBT Smart MALDI Biotyper (Bruker Daltonics, Bremen, Germany). We spotted all strains from UHRC in duplicate onto the MALDI target plate and overlaid with 1 μl of 70% formic acid. After drying at room temperature, we covered and dried the spots with 1 μl HCCA matrix, according to the manufacturer’s indications (Bruker Daltonics). We obtained two spectra in the range of 2,000-20,000 Da on each spot, resulting in 4 spectra per isolate. The isolates from UHB were analysed in one spot per strain and one spectrum from spot was acquired.

### Data processing of MALDI-TOF MS protein spectra and development of predictive models

For both, feasibility, and supervised studies, we processed all MALDI-TOF MS spectral profiles with the Clover MS Data Analysis software (Clover Biosoft, Granada, Spain). We applied pre-processing pipeline to all protein spectra that consisted of: 1) smoothing -Savitzky-Golay Filter: window length=11; polynomial order=3- and baseline subtraction -Top-Hat filter method with factor=0.02-; 2) creation of an average spectrum per isolate; 3) alignment of the average spectra from different isolates -shift: medium; constant tolerance: 2 Da; linear mass tolerance: 600 ppm-; 4) normalization by Total Ion Current (TIC).

#### Unsupervised feasibility study

To study the feasibility of MALDI-TOF MS for differentiation of ECC species, we proposed an unsupervised study based on Principal Component Analysis (PCA), and t-distributed Stochastic Neighbour Embedding (t-SNE). For this purpose, an oversampled balanced dataset of each ECC species was used. We included in total, 126 spectra from the 7 ECC species analysed in this study (sourcing from UHRC and UHB), as indicated in **Table 1**.

**Table 1.**
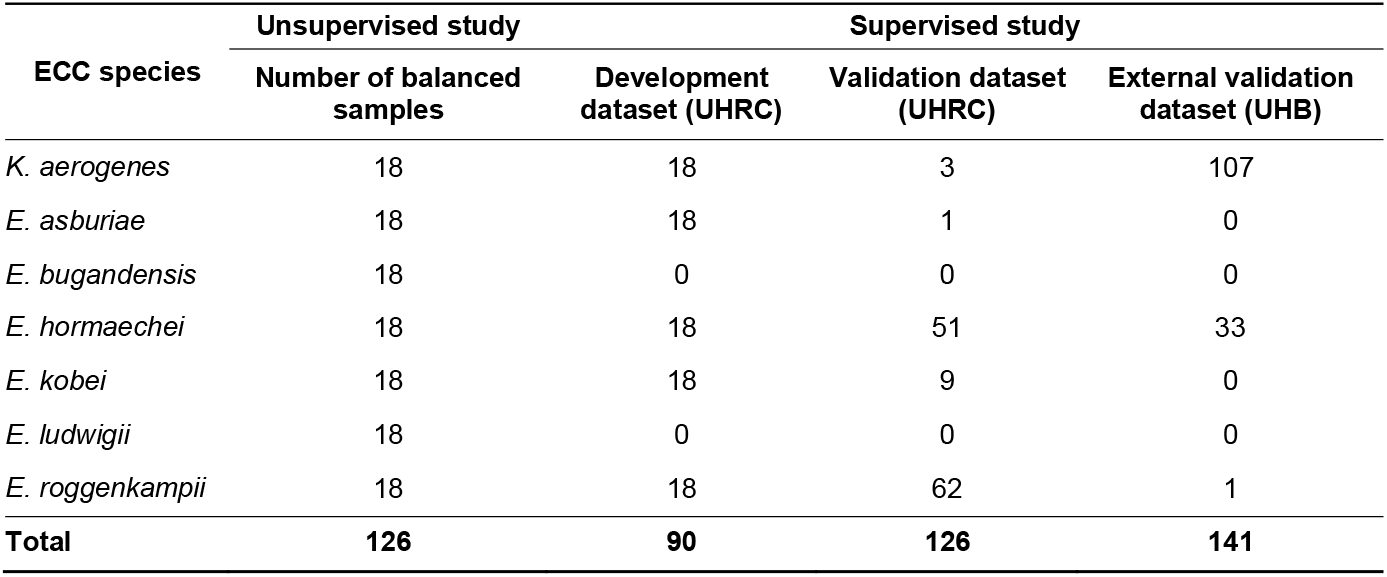
Number of ECC isolates used for the unsupervised feasibility study and the supervised model development.

#### Supervised model development

Once the feasibility of the study was determined, we proposed the supervised model development. In this case, three different datasets were created: training validation set, internal validation set, and external validation set. The details of these datasets are shown in **Table 1**.

Due to the lack of validation samples of *E. ludwigii* and *E. bugandensis*, these were not included in the development of the supervised model. Therefore, our supervised model was developed to predict the five ECC species: *E. asburiae* (cluster I), *E. kobei* (cluster II), and *E. hormaechei* (clusters III, VI, and VIII considered together), and *E. roggenkampii* (cluster IV). We applied three different supervised models: Partial Least Square-Discriminant Analysis (PLS-DA), Support Vector Machine (SVM) with Linear kernel (SVM-L) and with Radial Basis Function (SVM-R) kernel, and Random Forest (RF). The hyperparameter selection was performed by a 5-fold cross-validation technique.

Finally, we performed two external validations of the predictive models. First, 126 MALDI-TOF MS from UHRC and then 141 MALDI-TOF MS from UHB were blindly classified by the same predictive models using Clover BioSoft v0.6.1. This software uses the scikit-learn 0.23.2 python library to implement all statistical methods used in this study. For reproducibility purposes under FAIR principles, free access to all spectra and to reproduce the analyses in this work can be found at the following url: https://platform.clovermsdataanalysis.com/public-repository.

#### MSI Database

Recently, an online database has been developed for the rapid differentiation of ECC species based on their MALDI-TOF MS protein profile (19). This database has free access (https://msi.happy-dev.fr/) and has been built using protein spectra from 42 ECC isolates characterized by sequencing the *hsp60* gene. This identification method is considered the state-of-the-art method for the identification of ECC isolates at the species level. Therefore, both external validation datasets were also identified using the MSI database as a comparison to the methods proposed in this article. As stated above, MALDI-TOF MS spectra associated to this study have been also made publicly available.

### Ethics statement

The Ethics Committee of the Gregorio Marañón Hospital (CEIm) evaluated this project and considered that all the conditions for waiving informed consent were met since the study was conducted with microbiological samples and not with human products. At the University Hospital Basel only anonymized data was used with the purpose of quality control and assay validation. According to the Swiss Human Research Act no specific consent is required in this case. Data was either acquired in routine microbiological diagnostics (excluding cases with a rejected general consent) or used from a previously published dataset (DRIAMS).

## RESULTS

### Feasibility study

To prove the feasibility of MALDI-TOF MS to differentiate ECC species, an unsupervised hierarchical clustering with PCA and t-SNE pre-processing was performed (**Figure 1**). The protein spectra of the seven ECC species (*E. asburiae, E. kobei, E. hormaechei, E. roggenkampii, E. ludwigii, E. bugandensis*, and *K. aerogenes*), which are equally represented in the model, were compared. The dendrogram built with these data showed three main clusters: one containing *E. hormaechei*, a second cluster with *K. aerogenes*, and a third cluster with the rest of the species. Inside the latter cluster, *E. bugandensis* strains were clustered together and so did *E. asburiae, E. ludwigii, E. kobei*, and *E. roggenkampii*, although in these four cases some of the spectra were clustered with the wrong species (**Figure 1A, 1B and 1C**).

**Figure 1.**
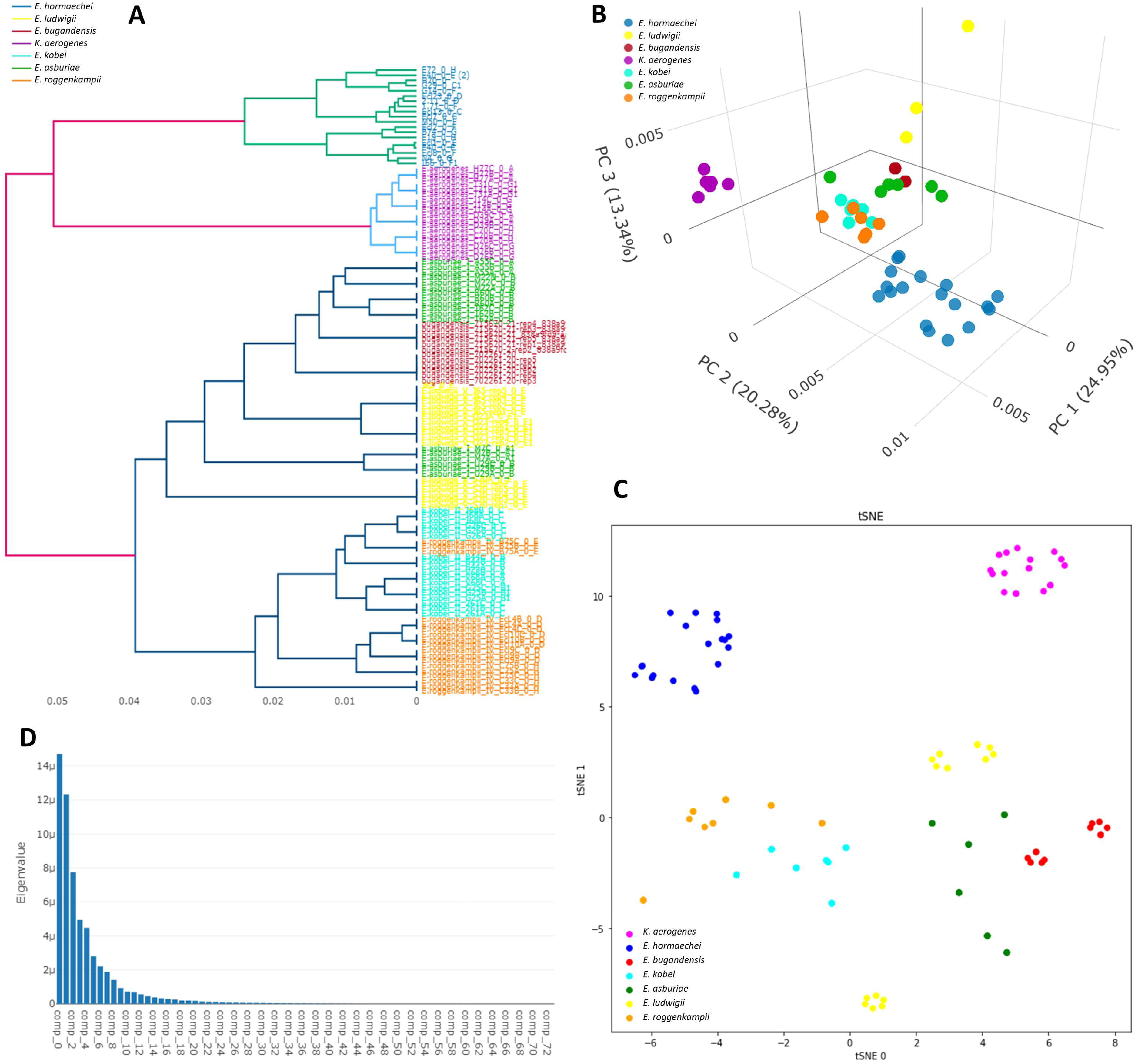
**A**. Dendrogram built with 126 MALDI-TOF MS spectra. **B**. PCA of feasibility study spectra. **C.** t-SNE of study spectra. **D**. PCA Eigenvalues showing the variance of each component.

The implementation of PCA to reduce the dimensionality showed that 14 components were needed to explain 95% of the variance (**Figure 1D**). This fact and the relatively accurate classification of ECC species using an unsupervised algorithm demonstrated the potentiality of MALDI-TOF MS to differentiate ECC species.

### Supervised models based on MALDI-TOF MS

To solve the limitations of unsupervised learning, we added the label knowledge to the training phase by using supervised algorithms such as PLS-DA, SVM, and RF. We trained these models using the development dataset shown in **Table 1**, and selected their hyperparameters by a 5-fold cross-validation technique. This cross-validation process led to the next hyperparameter selection: for PLS-DA 2 components were used, for SVM-L the value of C was 10, and for SVM-R the value of C was 10 and the value of γ was 1000. **Table 2** shows the results obtained for the internal 5-fold cross-validation, which have been further detailed in **Table S1**.

**Table 2.** Accuracy results for internal 5-fold cross-validation over development dataset (90 spectral profiles). PLS-DA: Partial Least Squares-Discriminant Analysis; SVM-L: Support Vector Machine-Linear kernel; SVM-R: Support Vector Machine-Radial Basis Function kernel; RF: Random Forest.

| Algorithm | *E. asburiae* |  | *E. hormaechei* |  | *E. kobei* |  | *E. roggenkampii* |  | *K. aerogenes* |  |
| --- | --- | --- | --- | --- | --- | --- | --- | --- | --- | --- |
| **PLS-DA** | 9/18 | 50% | 18/18 | 100% | 5/18 | 27.8% | 4/18 | 22.2% | 18/18 | 100% |
| **SVM-L** | 6/18 | 33.3% | 18/18 | 100% | 3/18 | 16.7% | 6/18 | 33.3% | 14/18 | 77.8% |
| **SVM-R** | **18/18** | **100%** | **18/18** | **100%** | **18/18** | **100%** | **18/18** | **100%** | **18/18** | **100%** |
| **RF** | **18/18** | **100%** | **18/18** | **100%** | **18/18** | **100%** | **18/18** | **100%** | **18/18** | **100%** |

Both *E. hormaechei* and *K. aerogenes* presented the same trend as in the feasibility study and their differentiation was 100% using non-linear approaches (SVM-R and RF) Only the implementation of a linear approach (SVM-L) yielded lower results for *K. aerogenes* (**Table 2**). For the rest of the analysed ECC species, we also obtained 100% correct classification by the application of non-linear approaches (**Table 2**).

In **Figure 2**, the distance between samples calculated by the RF classifier is shown. We detected a unique cluster for each species. Due to the results presented in **Table 2**, only SVM-R and RF were considered for further analysis. **Table 3** shows the results of SVM-R and RF for the validation dataset collected at UHRC and UHB.

**Figure 2.**
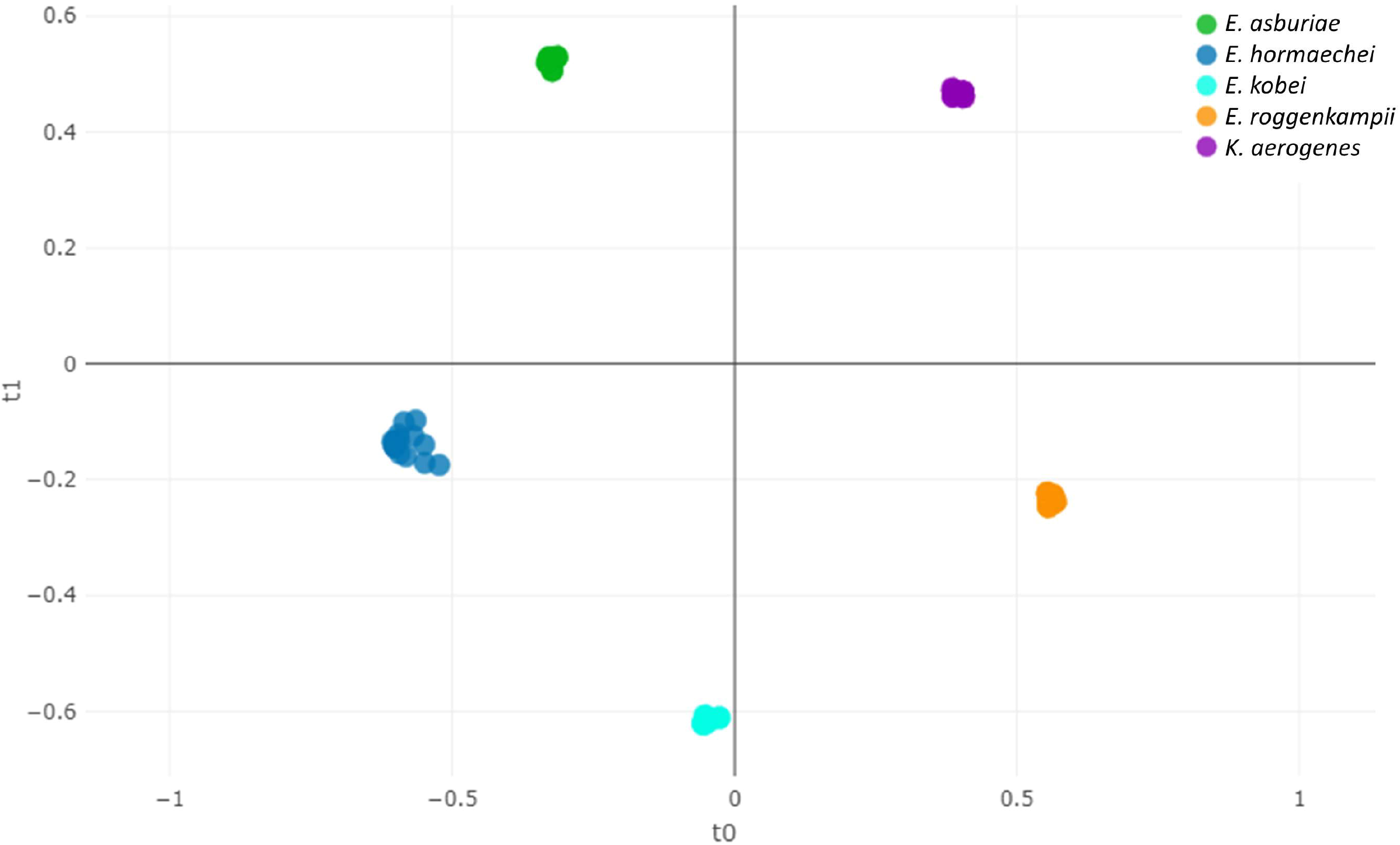
MALDI-TOF MS Euclidean distance between species by Random Forest classifier.

**Table 3.** Accuracy results for the validation dataset from UHRC and UHB, and the identification accuracy obtained by the MSI database. The MSI platform, specific for the identification of ECC species (19), was also applied for the classification of ECC species.

| Algorithm | *E. asburiae* |  | *E. hormaechei* |  | *E. kobei* |  | *E. roggenkampii* |  | *K. aerogenes* |  |
| --- | --- | --- | --- | --- | --- | --- | --- | --- | --- | --- |
| UHRC |  |  |  |  |  |  |  |  |  |  |
| SVM-R | 1/1 | 100% | 50/51 | 98.0% | 9/9 | 100% | 59/62 | 95.2% | 3/3 | 100% |
| RF | 1/1 | 100% | 50/51 | 98.0% | 9/9 | 100% | 59/62 | 95.2% | 3/3 | 100% |
| MSI | 1/1 | 100% | 50/51 | 98.0% | 9/9 | 100% | 60/62 | 96.8% | - | - |
| UHB |  |  |  |  |  |  |  |  |  |  |
| SVM-R | - | - | 15/33 | 44.1% | - | - | 1/1 | 100% | 102/107 | 95.3% |
| RF | - | - | 31/33 | 91.2% | - | - | 1/1 | 100% | 105/107 | 98.1% |
| MSI | - | - | 28/33 | 84.8% | - | - | 1/1 | 100% | - | - |

Both algorithms, SVM-R and RF, yielded the same results in the external validation performed on the MALDI-TOF MS spectra from the validation dataset sourcing from the same hospital (UHRC). In this case, all *K. aerogenes, E. asburiae*, and *E. kobei* isolates were correctly classified meanwhile one *E. hormaechei* strain was misclassified as *E. kobei* with both algorithms. For *E. roggenkampii*, two isolates were misclassified as *E. hormeachei* and one as *E. kobei* (**Table S2**). The accuracy of the model is shown in **Figures 3A and 3B**.

**Figure 3.**
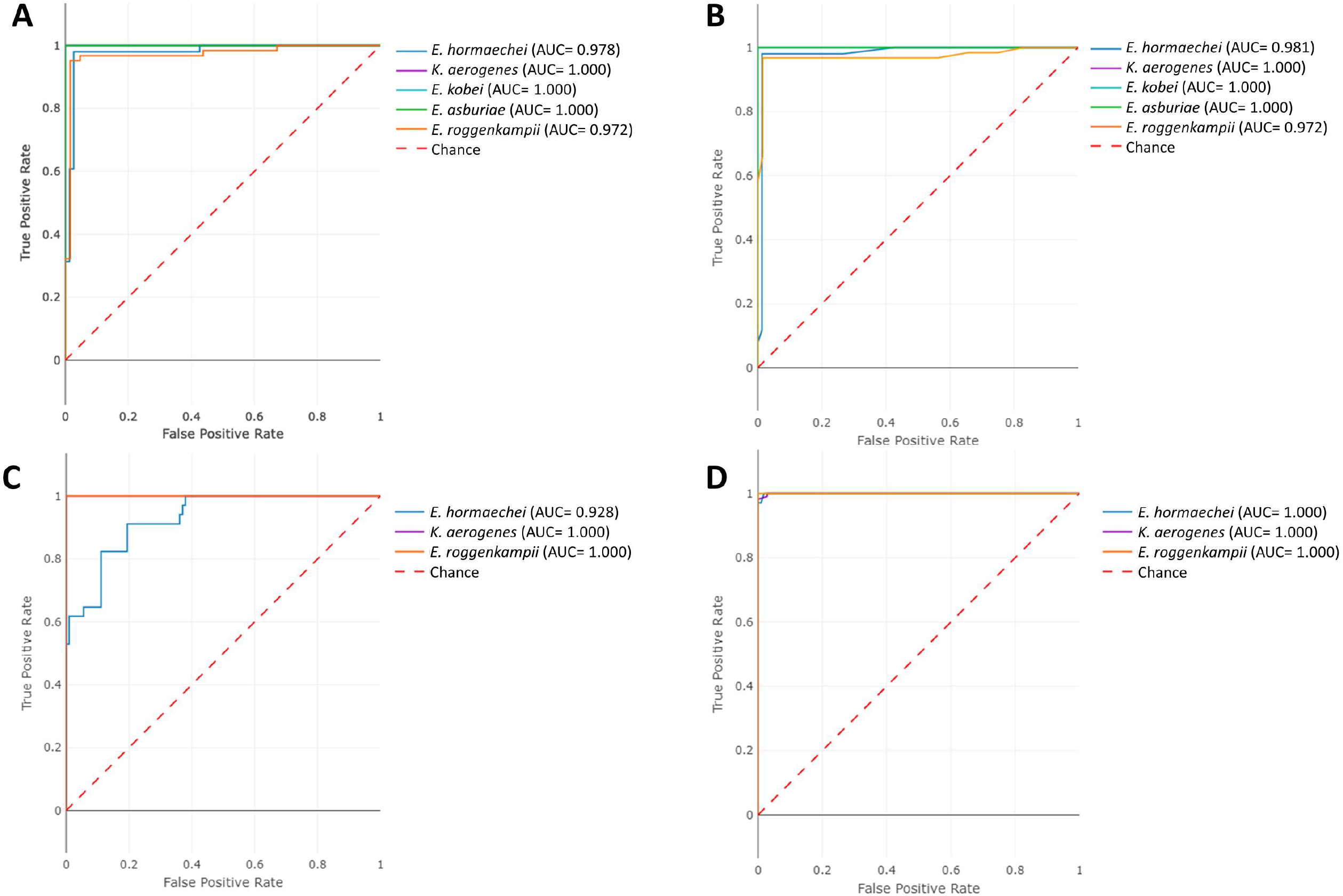
ROC Curves and AUC values for both SVM-R (A and C) and RF models (B and D) were applied to the external validations with MALDI-TOF MS protein spectra from UHRC (A and B) and UHB (C and D).

Since SVM-R and RF algorithms performed equally, both of them were considered for external validation with MALDI-TOF MS spectral profiles obtained at the UHB. In this case, 91.2% of the *E. hormaechei* (n=33), 100% of the *E. roggenkampii* (n=1), and 98.1% of the *K. aerogenes* isolates were correctly classified by RF, as shown in **Table 3**. The application of SVM-R yielded lower results for *E. hormaechei* and *K. aerogenes*. **Figures 3C and 3C** show the accuracy of both SVM-R and RF for the external validation collection from UHB.

### Identification of the ECC isolates using the MSI Database

Finally, the MSI Platform was also used as an identification tool for ECC species to compare the automatic approach developed in this study versus the current state-of-the-art method (19). Among the UHRC isolates, the identification rate for *E. asburiae, E. hormaechei*, and *E. kobei* was similar to the rates yielded by the predictive models developed in this study (**Table 3**). The only difference detected between both methods was that the MSI database correctly classified one more *E. roggenkampii* isolate. As for the protein spectra sourcing from the UHB, 84.8% of the *E. hormaechei* and 100% of the *E. roggenkampii* isolates (n=1) were correctly identified with the MSI platform. In this case, the MSI platform provided a lower rate of correct identifications for *E. hormaechei* than the RF algorithm (**Table 3**).

## DISCUSSION

In this study, the implementation of supervised, non-linear algorithms (SVM-R and RF) to MALDI-TOF MS spectra allowed the correct species assignment of 100% isolates belonging to two ECC species (*E. asburiae* and *E. kobei*) and between 91.2% and 98.1% for *E. hormaechei, E. roggenkampii*, and *K. aerogenes* (formerly *E. aerogenes*) sourcing from two different hospitals.

Poor discrimination of *E. cloacae* complex species by MALDI-TOF MS has been previously reported either by using commercial (11, 15) or enriched, in-house databases (20). However, a recent study from a research group with broad experience in MALDI-TOF MS and the creation of expanded libraries reported 92.0% correct species-level identification by implementing a specific in-house library enriched with well-characterized ECC strains and correct discrimination of 97.0% *E. hormaechei* isolates (19). This approach can be useful for the discrimination of close-related species, but the construction of a database is cumbersome and requires highly trained personnel. The implementation of the MSI platform allowed 94.9% correct species-level identification of 155 ECC protein spectra in this study. This rate was slightly lower than the obtained with the non-linear algorithms proposed by our approach.

In this study, we demonstrate the feasibility of MALDI-TOF MS to identify species within the ECC. First, hierarchical clustering showed that it is possible to differentiate between species using the information contained in MALDI-TOF MS as reported before (20). Secondly, a supervised study using machine learning algorithms yielded the correct classification of all ECC species. Therefore, different supervised classification algorithms were implemented to correctly provide species assignment of ECC species. The internal validation experiment demonstrated that non-linear approaches, such as SVM-R or RF, were needed to correctly identify all species. Both models perfectly classified all samples in internal cross-validation.

To further demonstrate that the model can perform in different scenarios with data different than the spectral profile used for model training, we performed two validation experiments. First, we carried out a validation with MADI-TOF MS protein spectra sourcing from UHRC. From a total number of samples of 116, both SVM-R and RF only misclassified four isolates, i.e., a 96.5% of accuracy in classifying species within the ECC was yielded. Secondly, we performed an external validation with MALDI-TOF MS sourcing from UHB to simulated a real-world scenario. These MALDI-TOF MS protein spectra originated in a different epidemiological scenario and were processed by operators from the UHB. In this case, SVM-R showed to be overfitted to the UHRC distribution, which was already pointed out by the value of γ value, scoring an 83.7% of accuracy. On the other hand, the current state-of-the-art tool -the MSI database-performed better than SVM-R with 94.9% of accuracy although it was not able to distinguish the *K. aerogenes* (19). However, RF outperformed both approaches with over 96.0% of accuracy in identifying the three species. Hence, it is demonstrated that supervised machine learning algorithms are feasible and, indeed, applicable in microbiology laboratories

One limitation of this study was the fact that all UHRC isolates were carbapenemase producing isolates, because this was the source of the previously analysed collection (15). However, the present study provides the first proof of concept for differentiating ECC species based on machine learning. For a definitive validation, improvement, and implementation of these predictive models, future studies will involve strains from a more diverse epidemiological and geographical origin and characteristics. Besides, not all analysed ECC species could be represented in the external validation dataset due to the lack of isolates from the species *E. ludwigii* and *E. bugandensis*.

The present study provides promising results for differentiating ECC species based on machine learning and MALDI-TOF MS protein spectra. It also highlights the facts that MALDI-TOF MS data should be linked to WGS data in order to allow future work and providing a reference standard. The MALDI-TOF MS and machine learning approach has been demonstrated to be a rapid and cost-effective method, suitable for correct species-level assignment of closely-related species, as in the case of ECC. The use of spectra analysis tools is becoming user-friendly and easy to apply and its use may provide species-level identification in a fast and inexpensive way. Once the model is validated with a comprehensive number of ECC species, an open web application will be deployed to be used by the community freely.

## ACKNOWLEDGMENTS

This work was supported by the projects PI15/01073 and PI18/00997 from the Health Research Fund (FIS. Instituto de Salud Carlos III. Plan Nacional de I+D+I 2013-2016) of the Carlos III Health Institute (ISCIII, Madrid, Spain) partially financed by the European Regional Development Fund (FEDER) ‘A way of making Europe’. MM was funded by the Community of Madrid (Programa de Garantía Juvenil, PEJD-2017-PRE/BMD-5106) and DRT with a postdoc contract from the Intramural Funding Program of the IiSGM. BRS (CPII19/00002) is a recipient of a Miguel Servet contract supported by the FIS. AGL was funded by a predoc contract from the Intramural Funding Program of the IiSGM.

## AUTHOR CONTRIBUTIONS

Ana Candela: experimental part, formal analysis, data collection, validation, visualization, writing – original draft preparation and review/editing. Alejandro Guerrero-López: formal analysis, data collection, validation, visualization, writing – original draft preparation and review/editing. Miriam Mateos and Alicia Gómez-Asenjo: experimental part, formal analysis, and data collection. Manuel J. Arroyo, Gema Méndez, and Luis Mancera: data analysis, validation, writing – original draft preparation and review/editing. Marta Hernández-García, Rosa del Campo: experimental part, formal analysis, and writing, submission of isolates, original draft preparation, and review/editing. Aline Cuénod and Adrian Egli: submission of isolates, original draft preparation, and review/editing. Juan de Dios Caballero, Laura Martínez-García, Desirée Gijón, María Isabel Morosini, Patricia Ruiz-Garbajosa, Rafael Cantón and Patricia Muñoz: validation, writing and review/editing. Emilia Cercenado: conceptualization, formal analysis, validation, writing, and review/editing. David Rodríguez-Temporal: conceptualization, formal analysis, validation, original draft preparation and review/editing; Belén Rodríguez-Sánchez: conceptualization, project administration, formal analysis, supervision, validation, visualization, original draft preparation and review/ editing.

