## Supplementary Tables 1 and 2 for "Automatic discrimination of species within the *Enterobacter cloacae* complex using MALDI-TOF Mass Spectrometry and supervised algorithms"

**Table S1.** Accuracy of internal 5-fold cross-validation over development collection for all algorithms tested.

| **Actual/Predicted** | ***E. asburiae*** | ***E. hormaechei*** | ***E. kobei*** | ***E. roggenkampii*** | ***K. aerogenes*** | **% correct** |
| --- | --- | --- | --- | --- | --- | --- |
| **PLS-DA** |  |  |  |  |  |  |
| *E. asburiae* | 9 | 0 | 4 | 2 | 3 | 50% |
| *E. hormaechei* | 0 | 18 | 0 | 0 | 0 | 100% |
| *E. kobei* | 6 | 0 | 5 | 7 | 0 | 27.8% |
| *E. roggenkampii* | 7 | 0 | 7 | 4 | 0 | 22.2% |
| *K. aerogenes* | 0 | 0 | 0 | 0 | 18 | 100% |
| **Total PLS-DA** |  |  |  |  |  | **60.4%** |
| **SVM-L** |  |  |  |  |  |  |
| *E. asburiae* | 6 | 4 | 2 | 3 | 3 | 33.3% |
| *E. hormaechei* | 0 | 18 | 0 | 0 | 0 | 100% |
| *E. kobei* | 6 | 3 | 3 | 6 | 0 | 16.7% |
| *E. roggenkampii* | 4 | 4 | 4 | 6 | 0 | 33.3% |
| *K. aerogenes* | 0 | 4 | 0 | 0 | 14 | 77.8% |
| **Total SVM-L** |  |  |  |  |  | **52.7%** |
| **SVM-R** |  |  |  |  |  |  |
| *E. asburiae* | 18 | 0 | 0 | 0 | 0 | 100% |
| *E. hormaechei* | 0 | 18 | 0 | 0 | 0 | 100% |
| *E. kobei* | 0 | 0 | 18 | 0 | 0 | 100% |
| *E. roggenkampii* | 0 | 0 | 0 | 18 | 0 | 100% |
| *K. aerogenes* | 0 | 0 | 0 | 0 | 18 | 100% |
| **Total SVM-R** |  |  |  |  |  | **100%** |
| **RF** |  |  |  |  |  |  |
| *E. asburiae* | 18 | 0 | 0 | 0 | 0 | 100% |
| *E. hormaechei* | 0 | 18 | 0 | 0 | 0 | 100% |
| *E. kobei* | 0 | 0 | 18 | 0 | 0 | 100% |
| *E. roggenkampii* | 0 | 0 | 0 | 18 | 0 | 100% |
| *K. aerogenes* | 0 | 0 | 0 | 0 | 18 | 100% |
| **Total RF** |  |  |  |  |  | **100%** |

**Table S2.** Accuracy of validation collection from UHRC, UHB strains and MSI database.

| **Actual/Predicted** | ***E. asburiae*** | ***E. hormaechei*** | ***E. kobei*** | ***E. roggenkampii*** | ***K. aerogenes*** | ***E. bugandensis*** | **% correct** |
| --- | --- | --- | --- | --- | --- | --- | --- |
| **SVM-R UHRC** |  |  |  |  |  |  |  |
| *E. asburiae* | 1 | 0 | 0 | 0 | 0 | - | 100% |
| *E. hormaechei* | 0 | 50 | 1 | 0 | 0 | - | 98% |
| *E. kobei* | 0 | 0 | 9 | 0 | 0 | - | 100% |
| *E. roggenkampii* | 0 | 2 | 1 | 59 | 0 | - | 95.2% |
| *K. aerogenes* | 0 | 0 | 0 | 0 | 3 | - | 100% |
| **Total SVM-R** |  |  |  |  |  |  | **96.8%** |
| **RF UHRC** |  |  |  |  |  |  |  |
| *E. asburiae* | 1 | 0 | 0 | 0 | 0 | - | 100% |
| *E. hormaechei* | 0 | 50 | 1 | 0 | 0 | - | 98% |
| *E. kobei* | 0 | 0 | 9 | 0 | 0 | - | 100% |
| *E. roggenkampii* | 0 | 2 | 1 | 59 | 0 | - | 95.2% |
| *K. aerogenes* | 0 | 0 | 0 | 0 | 3 | - | 100% |
| **Total RF** |  |  |  |  |  |  | **96.8%** |
| **SVM-R UHB** |  |  |  |  |  |  |  |
| *E. hormaechei* | 5 | 15 | 14 | 5 | 0 | - | 44.1% |
| *E. roggenkampii* | 0 | 0 | 0 | 1 | 0 | - | 100% |
| *K. aerogenes* | 5 | 0 | 0 | 0 | 102 | - | 95.3% |
| **Total SVM-R** |  |  |  |  |  |  | **83.1%** |
| **RF UHB** |  |  |  |  |  |  |  |
| *E. hormaechei* | 1 | 31 | 2 | 0 | 0 | - | 91.2% |
| *E. roggenkampii* | 0 | 0 | 0 | 1 | 0 | - | 100% |
| *K. aerogenes* | 0 | 0 | 2 | 0 | 105 | - | 98.1% |
| **Total RF** |  |  |  |  |  |  | **96.5%** |
| **MSI** |  |  |  |  |  |  |  |
| *E. asburiae* | 1 | 0 | 0 | 0 | 0 | 0 | 100% |
| *E. hormaechei* | 0 | 78 | 0 | 4 | 0 | 2 | 92.9% |
| *E. kobei* | 0 | 0 | 9 | 0 | 0 | 0 | 100% |
| *E. roggenkampii* | 0 | 2 | 0 | 61 | 0 | 0 | 96.8% |
| **Total MSI** |  |  |  |  |  |  | **94.9%** |
